# Unsupervised Machine learning to subtype Sepsis-Associated Acute Kidney Injury

**DOI:** 10.1101/447425

**Authors:** Kumardeep Chaudhary, Aine Duffy, Priti Poojary, Aparna Saha, Kinsuk Chauhan, Ron Do, Tielman Van Vleck, Steven G. Coca, Lili Chan, Girish N. Nadkarni

**Affiliations:** Division of Nephrology, Department of Medicine, Icahn School of Medicine at Mount Sinai, New York, NY; Department of Genetics and Genomic Sciences, Icahn School of Medicine at Mount Sinai, New York, NY

**Keywords:** AKI, machine learning, sepsis, subtypes, MIMIC

## Abstract

**Objective:** Acute kidney injury (AKI) is highly prevalent in critically ill patients with sepsis. Sepsis-associated AKI is a heterogeneous clinical entity, and, like many complex syndromes, is composed of distinct subtypes. We aimed to agnostically identify AKI subphenotypes using machine learning techniques and routinely collected data in electronic health records (EHRs).

**Design:** Cohort study utilizing the MIMIC-III Database.

**Setting:** ICUs from tertiary care hospital in the U.S.

**Patients:** Patients older than 18 years with sepsis and who developed AKI within 48 hours of ICU admission.

**Interventions:** Unsupervised machine learning utilizing all available vital signs and laboratory measurements.

**Measurements and Main Results:** We identified 1,865 patients with sepsis-associated AKI. Ten vital signs and 691 unique laboratory results were identified. After data processing and feature selection, 59 features, of which 28 were measures of intra-patient variability, remained for inclusion into an unsupervised machine-learning algorithm. We utilized k-means clustering with k ranging from 2 – 10; k=2 had the highest silhouette score (0.62). Cluster 1 had 1,358 patients while Cluster 2 had 507 patients. There were no significant differences between clusters on age, race or gender. We found significant differences in comorbidities and small but significant differences in several laboratory variables (hematocrit, bicarbonate, albumin) and vital signs (systolic blood pressure and heart rate). In-hospital mortality was higher in cluster 2 patients, 25% vs. 20%, p=0.008. Features with the largest differences between clusters included variability in basophil and eosinophil counts, alanine aminotransferase levels and creatine kinase values.

**Conclusions:** Utilizing routinely collected laboratory variables and vital signs in the EHR, we were able to identify two distinct subphenotypes of sepsis-associated AKI with different outcomes. Variability in laboratory variables, as opposed to their actual value, was more important for determination of subphenotypes. Our findings show the potential utility of unsupervised machine learning to better subtype AKI.

## Introduction

Acute kidney injury (AKI) occurs in up to a quarter of hospitalized patients and has been repeatedly shown to be associated with increased morbidity and mortality.(1–4) Sepsis is the most common cause of AKI in critically ill patients admitted to the intensive care units (ICU). It was initially thought that sepsis-associated AKI is due to systemic hypotension leading to a decrease in renal perfusion resulting in renal ischemia and acute tubular necrosis (ATN). However, there is growing evidence of different mechanisms of sepsis-associated AKI with potentially different clinical characteristics and outcomes.(5)

Thus, AKI is not likely a single clinical entity, but likely a clinical syndrome comprised of several different subtypes. However, there has not been any investigation into identifying these subphenotypes, which are all labeled sepsis-associated AKI. Electronic health records (EHRs) especially in the ICU setting, collect thousands of data points per individual patient. While there have been previous studies showing that machine learning approaches using EHR data identify distinct subtypes in chronic disease, there have not been any studies in acute disease.(6)

Our primary aim was to determine if we could identify subphenotypes of sepsis-associated AKI utilizing measurements done as part of patients’ routine care outside of traditional features such as age, gender, race, and comorbidities. We sought to incorporate hundreds of data features collected routinely in the EHR using unsupervised machine learning to identify subphenotypes of AKI in patients admitted to the intensive care unit with sepsis and explore differences in patient outcomes between the clusters.

## Methods

### Study Population

We utilized the Medical Information Mart for Intensive Care (MIMIC-III) database to identify patients with sepsis-induced AKI. MIMIC-III is a freely accessible critical care database of patients from a large, single center tertiary care hospital (Beth Israel Deaconess Medical Center in Boston, Massachusetts) from 2001 to 2012.(7) This database includes patient demographics, vital signs, laboratory results, billing codes, and notes. We included patients in the analyses if they had AKI within 48 hours of ICU admission as per Kidney Disease: Improving Global Outcomes (KDIGO) Guidelines.(8) We then defined sepsis with the Clinical Classification Software (CCS) which groups discharges into mutually exclusive categories utilizing International Classification of Diagnosis – Ninth Revision Codes (ICD-9).(9) We defined patient co-morbidities using the Elixhauser Comorbidity Software, which identifies comorbidities by grouping ICD-9-CM codes from hospital discharge records.(10) We excluded patients if they were less than 18 years old, admitted for ≤24 hours, end stage renal disease (ESRD), or missing vital signs. A study flow diagram is included in **Supplemental Figure S1**. As patients could be admitted several times during the 11 year period and develop AKI, we considered only the data from the first admission with AKI per person.

### Data Processing

We utilized laboratory values and vital sign measurements to identify the clusters and considered all labs and vitals from admission to 48 hours after the diagnosis of AKI for inclusion. We excluded data features which were missing in > 70% of patients. As there are fundamental differences in the measurements of the laboratory tests and vital signs features, they were processed separately. For vital signs, we had a feature space of 30 (including median, SD and count). The laboratory results feature space was much larger with 691 unique features with at least one value. We removed the median features with >70% missing values; corresponding SD and count were added later to the remaining lab features. For both class of features, we used k-nearest neighbor (knn)-based imputation method using 2 neighbors. *impute.knn* function was used in the *impute* R package.(11)

Since the laboratory results and vital signs have a different range of values, we used YeoJohnson (YJ) normalization separately to normalize the feature space.(12) For each class, features that were highly correlated (correlation coefficient >0.50) were excluded. The absolute values of pair-wise correlations were considered. If two variables have a high correlation, the function looks at the mean absolute correlation of each variable and removes the variable with the largest mean absolute correlation. This step was done to remove redundant features that added no additional information to the downstream clustering method. Subsequently, the features from labs and vitals were combined together and subjected to Box-Cox normalization to bring all features to a comparable scale.(12) Finally, the data were translated and transformed at log scale just prior to actual k-means clustering. Throughout the data processing steps, laboratory features derived from albumin, bicarbonate, and potassium were kept as we clinically adjudicated them to be meaningful.(13, 14) Selected features contributed to the final unsupervised clustering.

### Clustering

With the final feature matrix of combined labs and vitals for all the samples, we performed unsupervised clustering using k-means. We opted to generate 25 initial configurations for a range of cluster numbers (k=2 to k=10). We calculated the silhouette score in order to select the best cluster (k). Apart from that, we performed a 10-fold cross-validation using the features among the clusters corresponding to the final selected k. We used the random forest algorithm implemented in the *caret* packag*e*. We opted for 5 variables to be randomly sampled as candidates at each split and used 5 trees to grow in the 10-fold cross validation. Further, we picked each of the features and measured the difference in means between the two clusters. This was done to see the feature importance in each cluster with respect to each other. After obtaining the cluster labels, we used t-Distributed Stochastic Neighbor Embedding (tSNE) technique to reduce data to three dimensions for better visualization. We used the default parameters in the *Rtsne* package with perplexity=40.(15) Finally, clusters were visualized in 3D space using the *scatterplot3d* package in R.(16)

### Statistical Analysis

After cluster identification, we conducted analysis to explore differences between clusters. We used t-test for continuous variables, and Fisher’s exact test or chi-Square for categorical variables. We included need for renal replacement therapy, mechanical ventilation, and in-hospital mortality as outcomes of interest. We used log binomial regression to determine the association between cluster and adverse outcomes while accounting for patient characteristics that were significantly different on univariate analysis. We chose log binomial regression instead of logistic regression due to the high rates of in-hospital mortality. As this study was done on publically available, de-identified data, it was considered IRB exempt. Analysis was done using SAS 9.4 and R 3.4.3 software.

## Results

### Clinical Features of Patients with Sepsis-associated AKI

We identified 1,865 patients who had sepsis-associated AKI. Patients had a mean age of 66.3±15 years; 57% were men, and 75% were white. Most patients were admitted to the medical intensive care unit (MICU) (70%). Patients had a high prevalence of hypertension (51%), cardiac arrhythmias (38%), and diabetes mellitus (34%).

### Feature selection

Ten vital signs were available in the MIMIC-III database. Each vital sign was included as a median, SD, and count (number of times measured) for a total of 30 features. We identified 691 unique features corresponding to laboratory results, the total feature space including median, SD, and count was 2,073. After implementing data preprocessing (see Methods), total feature space after combining these vitals and labs contained 135 features. The feature selection step further reduced the number of features to 59 (9 vitals and 50 labs). Missingness of the original features and final features are presented in **Supplemental Figure 2** and **3** respectively.

### Unsupervised Clustering to identify Subphenotypes

From these combined features with transformed values, we implemented the k-means clustering ranging from k=2 to k=10. We relied on silhouette score for the independent assessment and selection of the cluster selection. We found k=2 with maximum silhouette score of 0.62; which was highest among the other cluster runs (**Supplementary Figure 4**). Cluster 1 had 1,358 patients while cluster 2 had 507 patients. **(Figure 1)** To assess robustness of clusters, we performed 10-fold cross-validation after we identified cluster labels. We obtained an average of 97.6% accuracy using the random forest algorithm.

**Figure 1:**
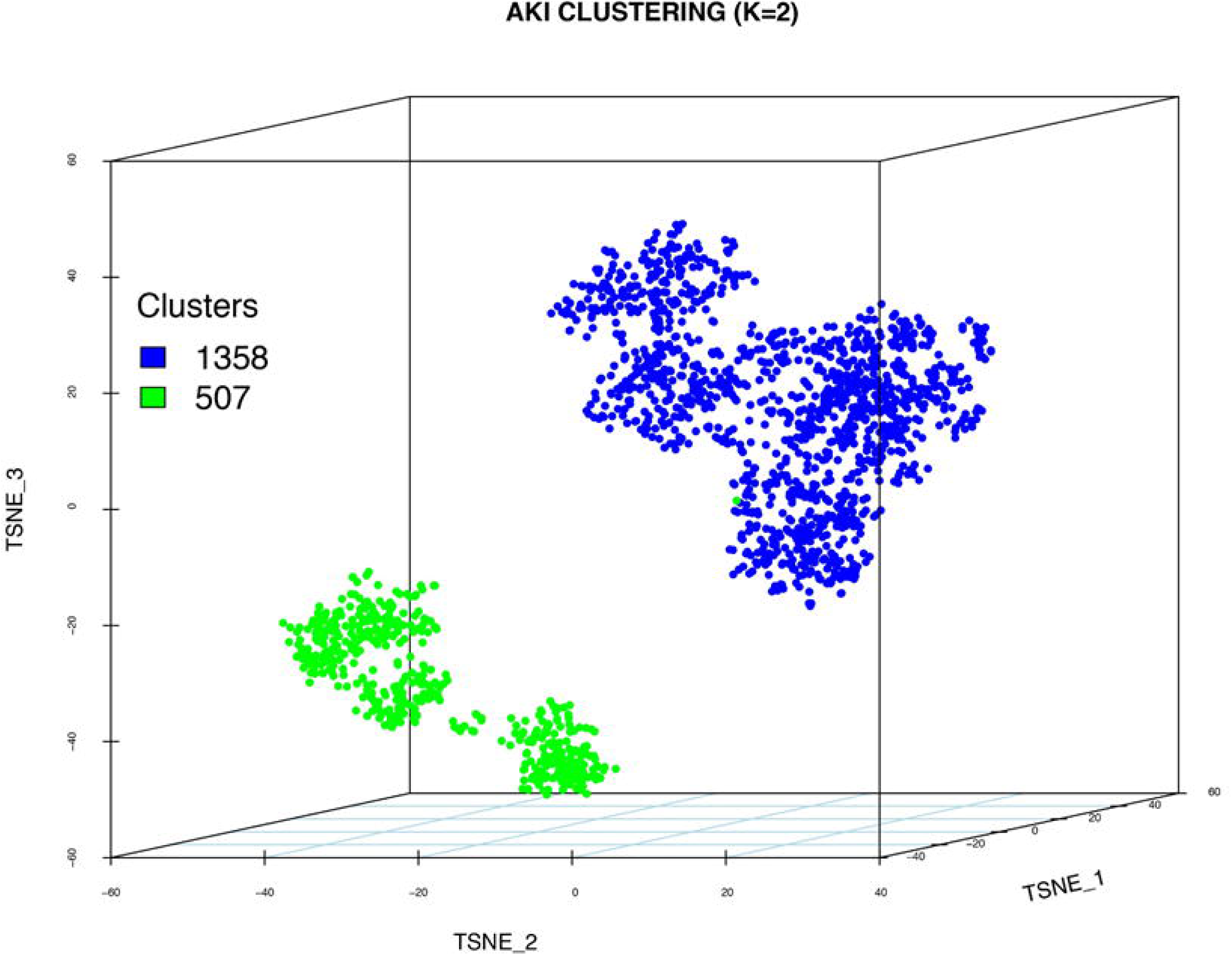
3-D Representation of clusters utilizing t-Distributed Stochastic Neighbor Embedding (t-SNE) method. This method condenses the 59 features into 3 transformed values which allows for 3-D representation. Each dot represents a single patient. The separation between two dots represents differences of features between two patients. Cluster 1 is represented in blue while Cluster 2 is represented in green.

### Clinical and Biological Characteristics of Each Phenotype

We identified several patient and admission characteristics that were significantly different between clusters. **(Supplemental Table 1)** Cluster 1 patients had higher prevalence of congestive heart failure (CHF) (37% vs. 31%, p=0.009) and lower prevalence of diabetes mellitus (DM) (36% vs. 39%, p=0.01). Mean Simplified Acute Physiology Score (SAPS) II scores were lower in Cluster 1, 45±15 vs. 47.6±16, p=0.002. Patients in cluster 1 were less likely to be admitted to the MICU (68% vs 76%, p=0.001) and less likely to have urgent/emergency admissions (97% vs 99%, P=0.02). There were statistically significant differences in several laboratory features such as hemoglobin, platelets, sodium, bicarbonate, and albumin however the absolute differences were relatively small. Cluster 1 had significantly higher systolic blood pressure (SBP), lower heart rate, and lower respiratory rate.

To determine the importance of each feature on clustering, we determined the mean difference between two clusters at the log scale (Figure 2). The largest differences were seen in basophil variability, eosinophil variability, creatine kinase and ALT variability. Features such as sodium, temperature, and pH level were not substantially different between clusters and likely had a smaller effect on cluster determination.

**Figure 2:**
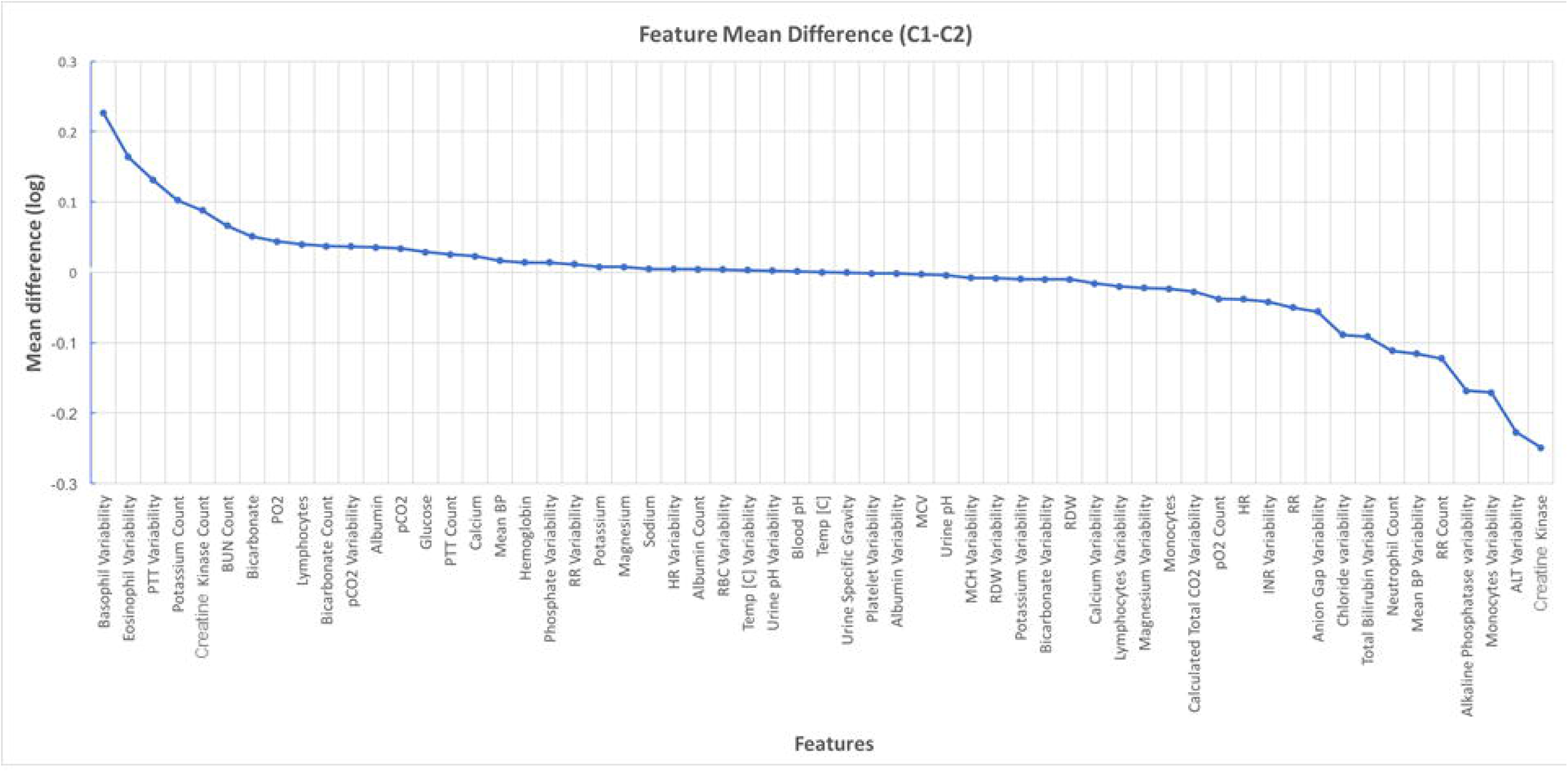
Mean Difference of Normalized Features between Cluster 1 and Cluster 2.

### Association between Phenotype and Outcomes

There was no difference in dialysis need between the two clusters (Table 1). However, Cluster 2 had lower proportion of patients requiring mechanical ventilation (49% vs. 54%, P=0.03) and higher rates of in-hospital mortality (25% vs. 20%, P=0.008). 30-day mortality was also higher in Cluster 2, however this was not statistically significant (30% vs 26%, P=0.11). Patients in Cluster 2 had a 40% higher risk of in-hospital mortality after adjustment for age, gender, ethnicity, CHF, DM, hematologic malignancy, and first ICU care unit with an adjusted odds ratio of 1.4, 95% CI 1.1-1.8.

**Table 1:** Differences in outcomes by cluster

|  | **Cluster 1**<br>**N=1358** | **Cluster 2**<br>**N=507** | **P value** |
| --- | --- | --- | --- |
| **Outcomes** |  |  |  |
| Any Dialysis | 134 (10) | 45 (9) | 0.52 |
| CRRT | 74 (6) | 29 (6) | 0.82 |
| CRRT duration (hours) | 126±115 | 109±94 | 0.5 |
| Mechanical Ventilation | 739 (54) | 248 (49) | 0.03 |
| Mechanical ventilation duration (hours) | 157±178 | 172±173 | 0.2 |
| In-hospital Mortality | 267 (20) | 129 (25) | 0.008 |
| 30-day Mortality | 354 (26) | 151 (30) | 0.11 |

## Discussion

We identified two distinct clusters of patients from patients within the larger syndrome of sepsis-associated AKI using unsupervised machine learning on routinely measured laboratory measurements and vital signs. Measures of variability were important to the identification of clusters. Clusters were significantly different in regards to comorbidities, laboratory measurements, and vital signs. We also found that these subphenotypes differed significantly in terms of mortality and mechanical ventilation.

There has been speculation that AKI in the ICU is not a single clinical entity but likely a complex syndrome comprising of several different subtypes.(17) Due to widespread use of EHRs, granular data are collected as part of routine clinical care on every ICU patient. These massive troves of data provide us with the opportunity to investigate this hypothesis in a data-driven manner. Subendophenotyping in chronic disease, such as diabetes has been conducted using similar EHR data with great success, and patient subgroups are found to have differing outcomes and genetic pathways.(6) Previous work, using trial data from an acute respiratory distress syndrome (ARDS) clinical trial has shown that there exist different subtypes with differing outcomes.(18) However, to the best of our knowledge this is the first instance of utilizing routinely collected EHR data from the ICU setting for subphenotyping of sepsis-associated AKI. Thus, this reinforces the concept, that even acute derangements may have distinct clinical types, an important implication for personalizing care to each individual.

We found that in-hospital mortality was 25% higher in patients in cluster 2. Cluster 2 patients had similar demographics and co-morbidities as Cluster 1 patients which indicate the clustering was not driven by these features. Additionally although features included in most ICU prediction models for mortality were also included as features in our clustering model, they likely played a small role in clustering as differences between the clusters on these variables were small.(19, 20) Of note, there was no difference in renal function parameters of KDIGO AKI stages, creatinine, or blood urea nitrogen (BUN) levels. Additionally, there was no difference in dialysis need or continuous renal replacement therapy (CRRT) need. This was surprising as there is abundant literature stating that severity of AKI, especially the need for dialysis, is associated with high morbidity and mortality.(21, 22) This may be partially explained by our time restriction of including labs only prior to AKI diagnosis and within 48 hours after AKI diagnosis.(23) These findings together highlight that individual features that are traditionally associated with differences in mortality in critically ill patients were not sufficient to identify the subphenotypes we have found here.

The purpose of this study was to agnostically identify subphenotypes within a larger clinical syndrome in a data driven manner. Thus, all laboratory features and vital signs were considered for potential inclusion into the analysis. The only limitations were to exclude features that were missing in >70% of patients (since they were unlikely to be informative) and highly correlated features. Additionally, we included measurements related to potassium, bicarbonate, and albumin into the model as these were considered clinically relevant features. We felt that it was not only important to include actual values but also the frequency and variability of continuous variables; and in fact 28 of the 59 features selected for inclusion were measures of variability and 9 of the 59 were counts. This is a significant advancement compared to previous models of clustering, where only summary measures of the predictors (mean/median) are used for either predictive modeling or clustering.

Through this inclusive data-driven approach, we included several features/factors that are not traditionally considered as related to ICU mortality into our clustering method including variability in key laboratory parameters. There is growing data that variability in values are important predictors of adverse events that should be considered in the clinical care of critically ill patients.(24–26) Eosinophil variability was among the factors with the largest difference between clusters. It has been established that eosinopenia occurs during an acute infection.(27) Several studies have found that eosinopenia can be used as a marker of sepsis on admission to MICU and is a significant predictor of ICU 28-day all-cause mortality.(28, 29) However, no studies have evaluated eosinophil variability and the association with outcomes in AKI. Creatine kinase was also higher in cluster 2 patients, who had higher mortality. Creatine kinase elevations can be seen in rhabdomyolysis, acute myocardial infarctions, and strokes. The etiology of creatine kinase measurements in this cohort of patients is currently unclear. However, it has been documented that hospitalized patients with fevers, especially those with bacteremia have been found to have elevated creatine kinase.(30) Additionally, observational data suggests an association between creatine kinase elevations and adverse outcomes. (31, 32) Thus, our data suggest, that creatine kinase may be an unexplored biomarker of risk in patients with septic AKI.

The results of our study should be considered in light of some limitations. We used the CCS category of septicemia to define our sepsis population; therefore we are unable to determine the timing of sepsis diagnosis and AKI diagnosis. However, this was mitigated by limiting AKI diagnosis to within 48 hours of ICU admissions and thus identifying patients with sepsis-associated AKI at the expense of a smaller sample size. We only included laboratory results and vital signs into our clustering algorithm. We decided not to include demographics and comorbidities, since we wanted to explore whether unbiased biochemical and biometric measurements could lead to viable clustering approaches. Indeed, the absolute difference in clusters on the basis of demographics and comorbidities was small and did not explain the difference in mortality. As to be expected, a majority of patients in both clusters was admitted to the MICU as their first ICU service. However, there was a notable difference between clusters on ICU first service, with more Cluster 2 patients being admitted to the MICU. There are inherent differences between patients admitted to different specialty units on admission diagnoses, nosocomial infection rates, and mortality. (33–35) Unfortunately, we are unable to identify medical patients who were boarded in non-medical ICUs as this may have an impact on mortality. Finally, while the MIMIC-III database is a large, granular, ICU database; it is a single center database. Although, we did cross-fold validation to ensure the clusters were robust, external validation in ICU databases from different centers is needed.

In conclusion, we were able to identify two distinct subphenotypes of sepsis-associated AKI using multidimensional biochemical and biometric data. These subphenotypes had similar baseline demographics, comorbidities, and AKI severity; however Cluster 2 patients had worse in-hospital mortality. Several factors which are not classically associated with adverse outcomes in sepsis-induced AKI were important contributors to the identification of subphenotypes. This approach could serve as a first step to identify clinical subtypes within the septic AKI syndrome and when combined with other –omics data, could help identify dysregulated pathways which could be targeted for therapeutic intervention.

## Acknowledgements

None

